# Partial characterization and the evaluation of antimicrobial activities of silver nanoparticles biosynthesized from *Cymbopogon citratus* leaf extract

**DOI:** 10.1101/2021.11.01.466798

**Authors:** S M Rakib-Uz-Zaman, Mohammed Nimeree Muntasir, Ehsanul Hoque Apu, Sadrina Afrin Mowna, Mst Gitika Khanom, Shah Saif Jahan, Nahid Akter, M. Azizur R. Khan, Nadia Sultana Shuborna, Shahriar Mohd Shams, Kashmery Khan

## Abstract

**Background:** Silver nanoparticles (AgNPs) are toxic to microorganisms and can potentially kill multidrug-resistant bacteria. Nanoparticles can be synthesized in many different ways, such as physical or chemical methods. Recently, it has been found that plant molecules can perform the same reduction reactions necessary for the production of nanoparticles but in a much more efficient way.

**Results:** Here, green chemistry was employed to synthesize silver nanoparticles using leaf extracts of *Cymbopogon citratus*. Effects of different parameters such as temperature, pH and volume of plant extract were also tested using their absorbance pattern at different wavelengths. The Surface Plasmon Resonance (SPR) changed with the changes in parameters. Changes in temperature from 20°C to 60°C have changed the highest absorbance from 0.972 to 3.893 with an SPR of 470 nm. At higher pH (11.1), the particles become highly unstable and have irregular shapes and sizes. At lower pH (3.97), the peak shifts to the right, indicating a smaller but unstable compound. We have also investigated the effect of the volume of plant extracts on the reaction time. The sample with the highest amount of plant extract showed the most absorbance with a value of 0.963 at λ_max,_ which was calculated to be 470 nm. The total formation of the AgNPs was observed visually with a color change from yellow to brownish-black. UV-visible spectroscopy was used to monitor the quantitative formation of silver nanoparticles, showing a signature peak in absorbance between 400 and 500 nm. We have estimated the size of the nanoparticles as 47nm by comparing the experimental data with the theoretical value using Mieplot. The biosynthesized silver nanoparticles showed enhanced antibacterial activity against several multidrug-resistant bacteria, determined based on the minimal inhibitory concentration and zone of inhibition.

**Conclusion:** The findings of this study indicate that an aqueous extract of *C. citratus* can synthesize silver nanoparticles when silver nitrate is used as a precursor and silver nanoparticles act as antimicrobial properties enhancers. These findings can influence further studies in this field to better understand the properties and applications of nanoparticles.

## 1. Background

Particles having a diameter of less than 100 nm are termed as nanoparticles. Nanoparticles exhibit novel and improved properties based on specific features such as better size, distribution, and morphology when compared to their constituent larger particles of the bulk materials. Due to their small size, nanoparticles have a greater surface-to-volume ratio. The specific surface area of silver nanoparticles is essential for their catalytic activity and other associated features such as antibacterial properties [1,2].

For almost 2000 years, the medicinal benefits of silver have been recognized. Silver compounds have been employed in several antibacterial applications since the nineteenth century. Silver ions and silver-based compounds are widely acknowledged to be very lethal to microbes, including some of the most common bacterial strains. These properties make it ideal for a variety of purposes in the medical sectors [3]. Topical lotions and creams comprising silver are used to reduce burn infection and sores. Medical equipment and implants made from silver embedded polymers are another prominent application. Furthermore, silver-containing consumer products like colloidal silver cream and silver-integrated fibers are being utlized in sports and athletic equipment [4]. Due to their enhanced antimicrobial activity and their potential application in treating cancers, many researchers are now focusing their research in developing an effective way to synthesize the silver nanoparticles [5].

### 1.1 Mechanism of Silver nanoparticles

The exact mechanism that is responsible for the antimicrobial effect of silver nanoparticles is still not clear. A number of theories have been put forward to explain the antimicrobial effect of silver nanoparticles. Silver nanoparticles can attach to the cell wall of bacteria and penetrate it, which subsequently leads to the structural damage in the cell wall and membrane with altered permeability of the cell membrane. The nanoparticles are accumulated on the cell surface by forming ‘pits’. Addionally, the formation of free radicals by the silver nanoparticles may be also responsible for the cell death. Numerous studies suggest that, silver nanoparticles can form free radicals when in contact with the bacteria and can damage the bacterial cell membrane by forming pores which results in cell death [6]. It was also suggested that nanoparticles might also produce silver ions [7]. These ions can intereact and inactivate the thiol groups of many important enzymes [8]. Moreover, the inactivation of the respiratory enzymes by silver ions can also generate reactive oxygen species, which can also attack the bacterial cell.

### 1.2 Difficulties in nanoparticle synthesis

There are many physical, chemical, and biological methods depicted in various literature on how to synthesize silver nanoparticles. The chemical processes include numerous methods that use toxic substances or are expensive and therefore are the ‘not so favored’ synthesis methods [9]. The physical methods include a large number of high-end equipment, which are expensive and occupy a considerable amount of space. The synthesis of silver nanoparticles (AgNPs) with a tube furnace possess several disadvantages as a tube furnace is very lagre, requires a lot of energy (more than several kilowatts) while increasing the temperature around the source material and also need a great deal of time (more than 10 minutes) to be thermally stable [10]. The chemical methods include many toxic components that are harmful for human consumption, and they also require harsh physical and chemical conditions, which can be hazardous for human health. In addition, the toxic residues produced along with the nanoparticles during a chemical reduction process makes the AgNPs unusable for any kinds of biomedical application [11].

### 1.3 Synthesis of plant nanoparticles

There has been a significant interest in developing a new strategy to successfully synthesize silver nanoparticles eliminating the drawbacks of chemical and physical production methods. Considering this, the idea of green chemistry has attained considerable recognition, particularity the contept is mainly focused in replacing the use of harmful chemicals. Scientists are also focused on developing methods and technologies to reduce and where possible to completely eliminate the compounds harmful to human health and the environment [12]. Among several biological methods, the use of plant extracts for synthesizing AgNPs has gain immense popularity. Owing to its directness in procedures, ease of monitoring, easy sampling and lower costs, this plant-based technique can be adopted as a replacement to the regularly utilized chemical techniques to facilitate the widespread production of AgNP[12]. Plant-based nanoparticles are considered to be eco-friendly as their production methods can effectively replace chemical reduction processes [13]. The metabolites present in plant extracts may aid in the reduction process [13]. Moreover, plants are readily available, easy to grow and safe to handle.

Scientists are still investigating the mechanism by which plant extract can reduce the silver ions. Recently, Fourier transform infrared spectroscopy (FTIR) spectroscopy of biosynthesized AgNPs has revealed that the biomolecules present in plant extracts are responsible for synthesizing nanoparticles[14]. Terpenoids are one of the most biomolecules involved in this process. Terpenoids are commonly referred as isoprene, an organic molecule contains five-carbon isoprene compounds. Some studies have suggested that the *Geranium* leaf extract contains terpenoids which contribute the most in AgNPs biosynthesis[14]. Another major plant metabolite is flavonoids, a polyphenolic compound consists of 15 carbon atoms and are water-soluble[15]. That is why it is imperative that while synthesizing the nanoparticles from the plant extract, the plant must have a high amount of terpenoids and flavonoids and exhibit some medicinal properties of their own. In addition, genetic variations, environmental and ecological factors have made plants chemically very diverse which further facilitate their application in synthesizing AgNPs [16].

### 1.4 Selection of plant extract

*Cymbopogon*, also known as lemograss, belongs to the Gramineae family. This herb is very rich in essential oil content. *Cymbopogon* genus is commomly available in the tropical and subtropical areas of Asia, America and Africa. People have been using *Cymbopogon citratus* as everyday tea, insecticide, insect repellant and as medicinal supplement to treat anti-inflammatory and analgesic diseases for hundred of yeas all over the world. Medical application of this herb also includes cures for malaria and stomachache due to its antioxidant properties.

In the current study, this plant species was selected since it has phytochemicals such as flavonoids, alkaloids, tannins, carbohydrates, steroids, and phytosteriods in relatively high concentrations [17]. An aqueous extract prepared from the leaves of *Cymbopogon citratus* was used as both bioreductant and capping agent for the green synthesis of silver nanoparticles to study the effect of volume of plant extract, reaction temperature, and reaction pH on the silver nanoparticles’ stability, synthesis rate and particle size. The AgNPs were prepared using different volumes of *Cymbopogon citratus* extract, and the response was conducted under different physiological conditions and checked for quality using UV-Vis spectroscopy. Antibacterial effects of the synthesized nanoparticles were also examined by testing them against selected pathogens such as *E. coli* (ETEC), *Salmonella paratyphi, Bacillus cereus, Vibrio cholera, Shigella flexneri*.

### 1.5 Objectives of this study

The primary purpose of this study is to figure out if an aqueous extract of *Cymbopogon citratus* can synthesize silver nanoparticles when Silver nitrate is used as a precursor. Furthermore, different parameters will also be tested to optimize the reaction condition and determine which combination of physical and chemical conditions can provide the best results. Most importantly, since silver nanoparticles are potent antimicrobial agent enhancers, they will be tested for their antimicrobial properties and how they act when combined with the same plant extract used as the bioreactor for the reaction process.

## 2. Methods

### 2.1 Collection and Preparation of Plant extract

The whole plant of *Cymbopogon citratus is* collected from local nurseries and gardens. Formal identification of the plant was conducted based on its characteristics and available information from the nurseries. They were washed thoroughly with distilled water several times to remove dust and dried under shade. After drying, they were rewashed to remove any unwanted dust particles. Then the leaves were dried at room temperature to remove the water from the surface of the leaves. About 10 g of finely incised dried lemongrass leaves were boiled in 150 ml distilled water at 60 °C for about 10 minutes. The supernatant was filtered using Whatman filter paper No.1 to remove the particulate matter. A dark greenish yellow clear solution is obtained and stored at 4–8 °C.

### 2.2 Synthesis of silver nanoparticles with aqueous *C. citratus*

Two mM solution of silver nitrate was prepared by dissolving 0.017 gm of AgNO3 in 50 ml of distilled water as described in [5]. Six ml extract of lemongrass leaf was mixed with 34 ml of 2 mM AgNO3 solution. The effect of time was studied at regular intervals of 24, 48, 72, and 96 hours in the leaf extract. The development of pH was studied by adding NaOH to the solution and bringing the pH to about 11.1. Similarly, to make the solution more acidic, a few drops of concentrated HCL were added to make the pH of the solution 3.97. The reaction was also carried out at different temperatures, such as 20°C, 40°C, and 60°C. The mixture was stirred nonstop, maintaining aforementioned conditions for 15 min with a magnetic stirrer. The formation of silver nanoparticles by aqueous *C. citratus*. The silver nanoparticles were repeatedly centrifuged at 3000 rpm for 10 min. The resulting pellets were air-dried and then redisposed in deionized distilled water[18].

### 2.3 Characterization of the Silver Nanoparticles

#### 2.3.1 UV–visible spectrometric analysis

The UV−visible spectra of the synthesized silver nanoparticles were recorded as a function of wavelength using a UV−Vis spectrophotometer (Genesys 10s UV-Vis Spectrophotometer) operated at a resolution of 1 nm. The reduction of silver was measured periodically at 300–700 nm. A spectrum of silver nanoparticles was plotted with the wavelength on the x-axis and absorbance on the y-axis[18].

### 2.3.2 Estimating the size

The SPR of AgNPs with high symmetry, like spheres or ellipsoids, was calculated with reasonable accuracy by analytical expressions developed in the Mie theory frame [19]. The size was estimated with the software Mieplot (Version 4.6.13). The calculations were done by following the manual collected from online sources[20]. First, from the advanced drop-down menu, the refractive index was selected as a sphere, silver if particles present in the medium were spherical in nature. Then again, from the advanced drop-down menu refractive index, the surrounding medium, water was selected. To determine the particle, several calculations were performed and then compared with our experimental data. The particle size of the best-fit spectrum will be the particle size determined from Mie Theory [20].

### 2.4 Antibacterial Assays

#### 2.4.1 Bacterial Strain collection

The bacterial strains such as *E. coli* (ETEC), *Salmonella paratyphi, Bacillus cereus, Vibrio cholera, Shigella Flexneri* were collected from BRAC University laboratory stock. A loopful of the desired organism was streaked in standard NA (Nutrient Agar) media. The plates were then incubated at 37°C for 24 hours. Then they were stored for further use.

#### 2.4.2 Agar Well Diffusion method

A suspension of the organism to be tested is prepared in a saline solution and measured to be equal to 0.5 McFarland standard (1 × 10^8^ colony forming units (CFU)/ml). The cultures were swabbed on standard MHA (Mueller Hinton Agar) media with a sterile cotton swab. A well with a diameter of approximately 7-mm is made on a Mueller Hinton Agar plate with gel puncture. 60µl of synthesized particles, plant extract, plant extract combined with nanoparticles and AgNO_3_ solution was inoculated to the well. Then, the plates were incubated in an incubator at 37 °C for 24 h, and the zones of inhibition were discussed [21].

#### 2.4.3 Minimum Inhibitory Concentration

The bacterial cultures were grown in Nutrient Broth and transferred to sterile test tubes. Different concentrations of Silver nanoparticles (50µl, 100µl, 150µl, 200µl, 250µl) were added to the culture broth. The test tubes were incubated at 37°C for 24 h. After incubation, the growth of the bacterial isolates in the test tubes was observed as turbidity using a spectrophotometer at 600 nm. The least concentration where no turbidity was observed was determined and noted as the MIC value[22].

## 3 Results

### 3.1 Synthesis of Silver Nanoparticles

After the addition of plant extract to the silver nitrate solution, the solution started to change color. The solution was pale yellow when the reaction started. After 30 min of continuous stirring, the solution slowly started to turn darker. After 2 hours of reaction time, the solution turned completely dark brown, indicating the presence and formation of silver nanoparticles in the mixture.

### 3.2 Surface Plasmon Resonance Analysis

The silver nanoparticles do not show any absorbance below 390 nm. The highest absorbance was observed at 470 nm with an absorbance of 2.24 after 24 hours of reaction time. This highest peak is the Surface Plasmon Resonance (SPR) for the synthesized nanoparticles. After 470 nm, the absorbance slowly starts to drop as low as 0.5.

### 3.3 Analyzing the effect of temperature

Performing the reaction at different temperatures had a significant impact on the size and shape of the nanoparticles. The Surface Plasmon Resonance changed when the response changed temperature. At 20°C, the peak was observed at 471 nm, and the highest absorbance after two hours was 0.972. It also had a broader peak compared to the other spectrum. At 60°C, the peak absorbance was highest at 3.893 with an SPR of 470 nm. The shape of the spectrum was different from the others, with irregular and rigid shape towards the peak and the peak was also very well defined. The most well-defined peak was observed at 40°C temperature. While the curve was smooth, the peak was broader than the one with the highest temperature. The SPR was monitored at 469 nm, and the absorbance peaked at 2.237.

### 3.4 Effect of pH on the formation of Silver nanoparticles

The effects of pH on the reaction media can be observed with a visual inspection just by looking at the solution. Adding a few drops of concentrated HCL made the solution white. Letting the solution sit for a couple of hours made a white precipitate, which was volatile. The UV spectral analysis revealed that at a pH of (3.97) the peak was observed at 435 nm, which is a lot lower than the usual SPR of 470 nm. In the case of a higher pH (11.1), the solution quickly turned black with the addition of NaOH. The UV spectrum revealed a random absorbance pattern that did not fit the usual spectral pattern observed in nanoparticle synthesis.

### 3.5 Effect of Plant Extract Volume

The volume of the plant extract had a significant effect on the amount of nanoparticle production. Increasing the volume of the plant extract caused a significant difference in the absorbance pattern. During the 1-hour mark, the difference was minimal. As time progressed, during the 3-hour mark, the sample with the highest amount of plant extract showed the most absorbance with an absorbance of 0.963 at λ_max,_ which was calculated to be 470 nm. It was the same with the 4-hour mark, with the highest being the same with 12 ml of plant extract and absorbance of 2.031. In contrast, the lowest point was with 8 ml plant extract showing absorbance of 0.831.

### 3.6 Estimating the Size of the Silver Nanoparticles

Several calculations were made using the Mie Plot tool, each time using different nanoparticles as our standard model. It has been found that when the particle size is considered to 47 nm, the spectrum it shows in the theoretical model closely resembles the spectrum created from our experimental values, with. The theoretical model showing SPR position at 475 nm and the experimental model showing the SPR position at 470 nm. So, the size of nanoparticles using lemon grass plant extract can be considered to have a radius of approximately 47 nanometers.

### 3.7 Stability of the Silver Nanoparticles

The nanoparticles were stable for 96 hours without adding any sort of extra stabilizers. This is evident by observing the absorbance spectrum at different time intervals. At 24-hours, the absorbance peaked at 1.111 with the SPR positioned at 481 nm. Similarly, as time progressed, the absorbance increased. The maximum absorbance was observed at the 96-hour mark, where it showed an absorbance of 1.979. All four of the curves have a similar pattern with the SPR at 481 nm. The starting point was also identical in all four of these. Below 390 nm, all of them showed little to no absorbance. The absorbance increased as time progressed, and the peaks also became sharper and well defined.

### 3.8 Antibacterial Assays

#### 3.8.1 Results of Agar Well Diffusion

#### 3.8.2 Results of Minimum Inhibitory Concentration test

The minimum inhibitory concentration was calculated using standard broth dilution methods. It has been observed that *Salmonella paratyphi* requires the highest concentration of nanoparticles, which is 150 µl/ml. *Bacillus cereus* and *Shigella flexneri* required the lowest amount of 50 µl/ml of silver nanoparticles.

## 4.1 Discussion

The attempt to synthesize nanoparticles from aqueous *Cymbopogon citratus* was successful, as evident from (Figure 1). The color change indicates that nanoparticles were beginning to form in the reaction medium [5]. When a specific plant extract volume was added to the transparent color AgNO_3_ solution, it changed into a pale-yellow color. A while after the solution was allowed to sit at room temperature, it slowly began to change color. (Figure 1 (e)) shows that after 2 hours of reaction time, the solution became dark brown in color, which is one of the key indications that silver nanoparticles are present in the solution. The gradual color change can be observed from Figure 1 (a to e), which means that as time goes on, the concentration of nanoparticles in the solution was slowly increasing, as evident from the color change from yellow to a much darker tone of brown. After 24 hours of reaction time, no more change in color is observed, which means that all the silver molecules in the solution have already been reduced to silver nanoparticles.

**Figure 1:**
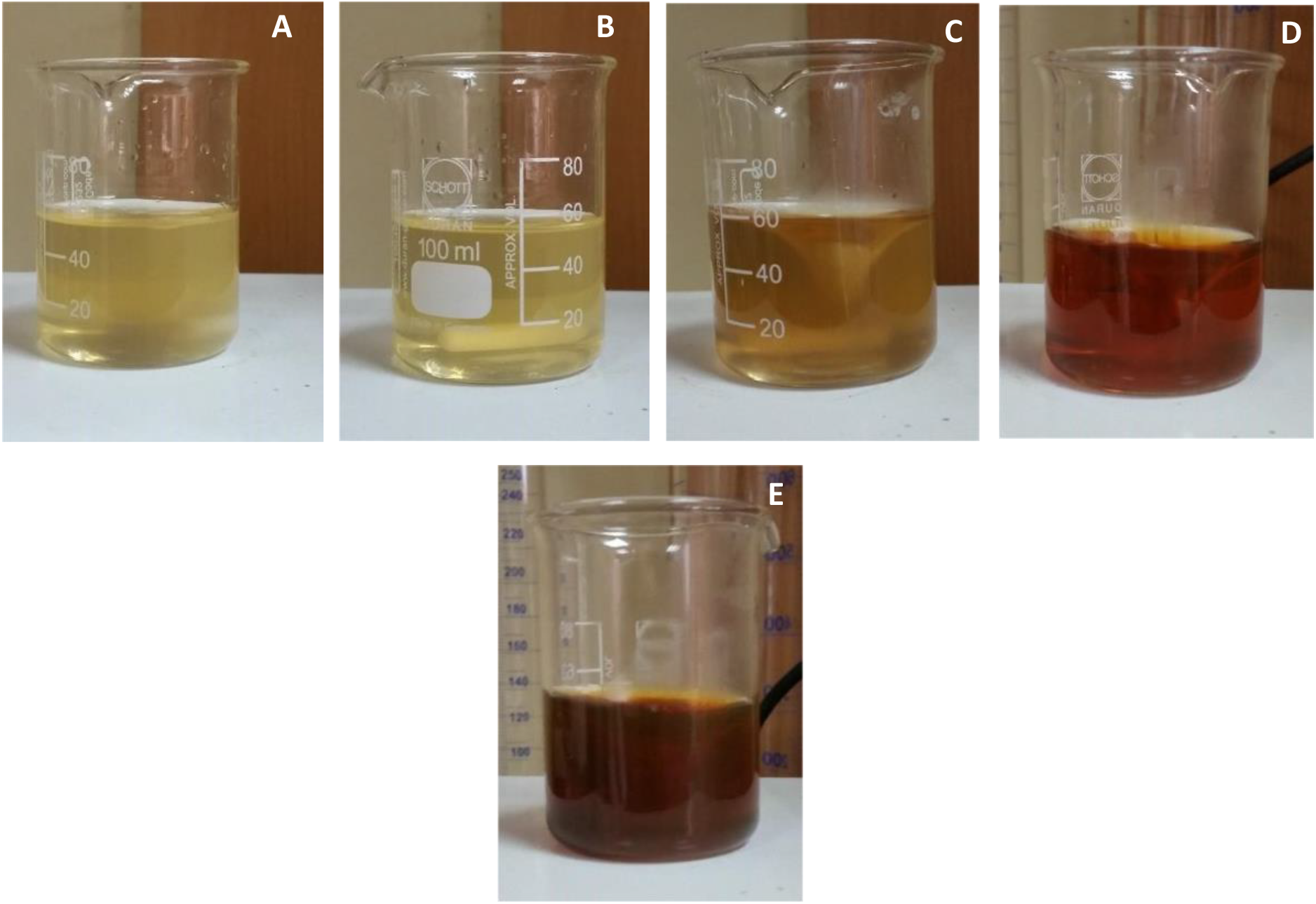
Synthesis of silver nanoparticles at different time intervals (a) 15 min, (b) 30 min, (c) 1 hour, (d) 1.5 hours, and (e) 2 hours. From (a) to (e), the colour of the solution turns from a pale yellow to colour to a dark brown colour. The colour is darkest at (e), indicating the presence of silver nanoparticles.

Even though the color change can indicate the presence of nanoparticles in the solution, they can only be confirmed by performing UV spectral analysis. Analyzing the absorbance peak, we can confirm whether silver nanoparticles are present or not. This is because the Surface Plasmon Resonance (SPR) of the silver nanoparticles determines their optical, physical, and chemical properties. In case of silver nanoparticles, the absorbance should peak between 400-500 nm[5,18]. This peak in absorbance is called their SPR value. Correlating the AgNPs plasmonic properties with their morphology is a fast and easy way to monitor the synthesis by UV–visible spectroscopy[23]. As we can see from Figure 2, the absorbance peaks at 470 nm, which means this is the SPR value of the nanoparticles we were able to synthesize. This peak is more extensive than our reference value[5], which might be because different plants work differently to reduce silver nanoparticles. That is why nanoparticles synthesized using diverse plant samples will have varieties in their shapes and sizes according to the models used as a bioreactor. Broader peaks indicate larger-sized particles, while smaller particles will form more distinct and well-defined peaks during the UV spectral analysis [19,24].

**Figure 2:**
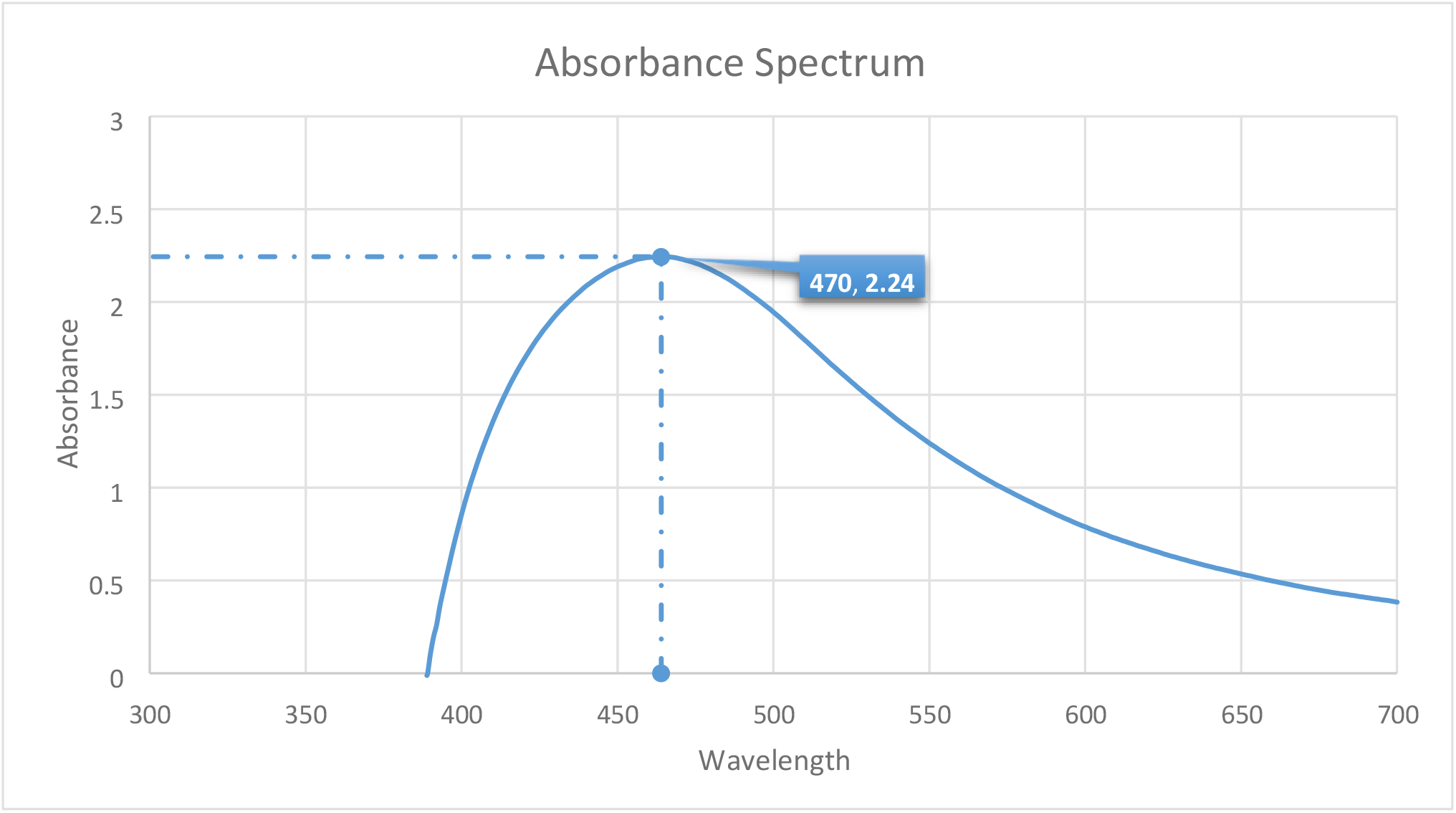
Absorbance Spectrum of synthesized Silver nanoparticles. The highest absorbance was observed at 470 nm with an absorbance of 2.24. After 470 nm, the absorbance slowly starts to drop, and the lowest absorbance recorded was 0.5 while the absorbance continues to go lower. There is no absorbance observed before an SPR of about 390.

The effect of temperature has an enormous impact on the size and shape of the newly formed silver nanoparticles. We can see clearly from Figure 3 how the spectrum changes concerning the temperature used in the reaction. As the temperature is increased, the peak slowly starts to rise and becomes narrower. At the highest temperature of 60°C, the peak became more defined, but the spectrum became uneven, and there was a lot of roughness in the spectrum. This was because the increase in temperature caused the nanoparticles to become highly unstable. The noise could also indicate that the solution was not uniform, and particles of various shapes and sizes were available inside the solution. The SPR spectra greatly vary with reaction temperature.

**Figure 3:**
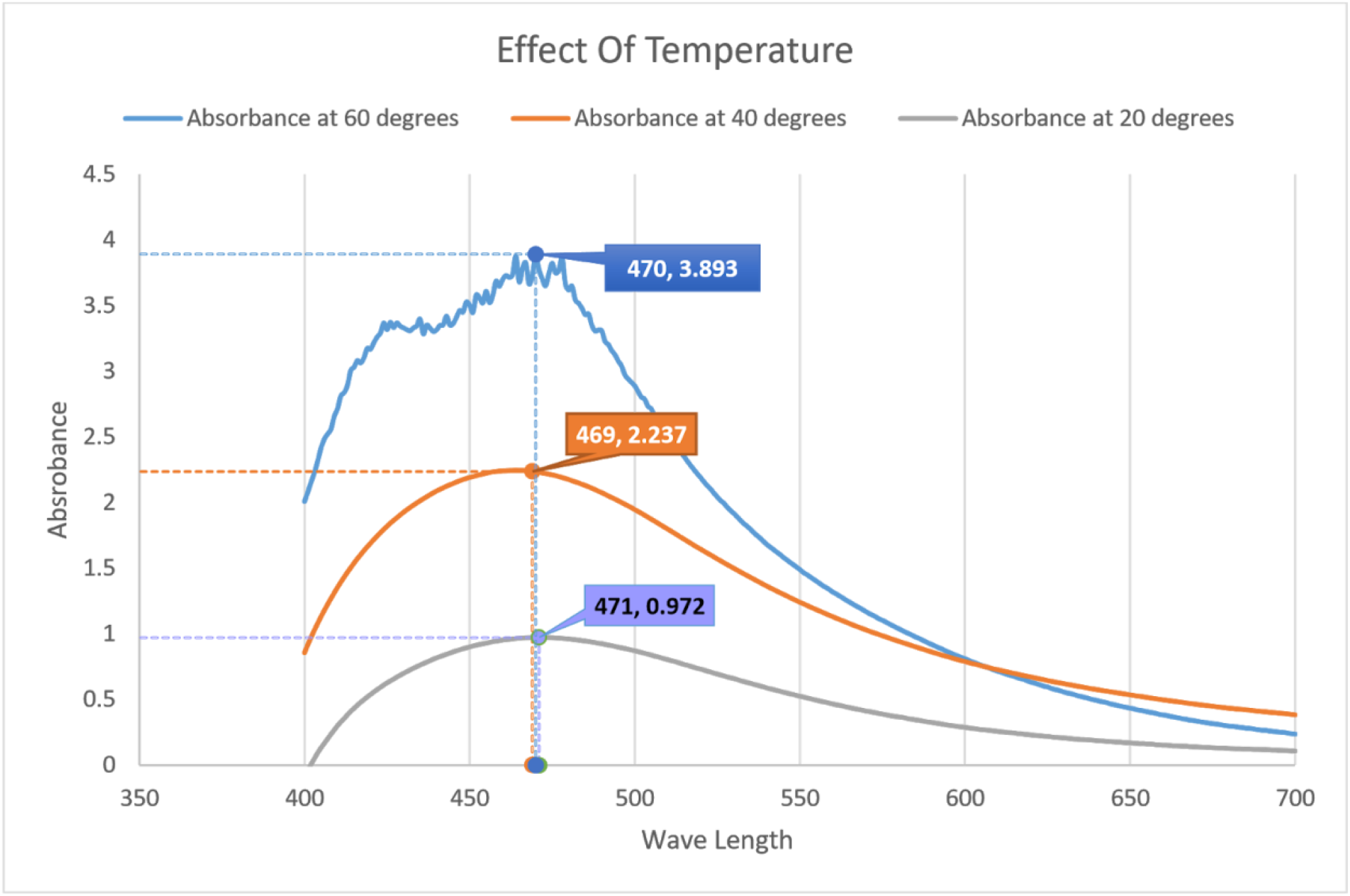
Effect of Temperature on the size and shape of nanoparticles. At 60°C, the peak absorbance was recorded highest at 3.893 with an SPR of 470 nm. At 40°C, the SPR was observed at 469 nm, and the absorbance peaked at 2.237. At 20°C, the peak was observed at 471 nm, and the highest absorbance 0.972

When reaction temperature increases, the absorbance peak increases with an increase in reaction rate, resulting in smaller sized nanoparticles[14,25]. The temperature enhances the rate of reduction, which results in decreased reaction time. Therefore, a rise in temperature may lead to smaller size AgNPs. The increase in temperature also increases the kinetic energy of molecules, which increases the consumption of silver ions. Thus, particle size growth and formation of uniform sized AgNPs is less likely[14,26].

The pH of the solution dramatically influences the size, shape, and optical properties of the nanoparticles. As evident from Figure 4, increasing the pH of the medium made the particles highly unstable, and we get a spectrum that indicates that. Lowering the pH to a more acidic medium made the solution completely white in an instant. Even though the spectrum is somewhat similar to our usual spectrum, we can notice that the peak shifted towards the right side and the size of the nanoparticles became much smaller. The nanoparticles in this medium were highly volatile and couldn’t be separated through repeated centrifugation as opposed to much heavier and more stable particles in the stable pH of 5.38. This could’ve happened for various reasons. The change in pH may lead to a change in charge of natural phytochemicals present in an extract. This charge change influenced the adherence of silver ions to biomolecules and might reduce silver ions to AgNPs. Due to the positive charge on silver ions the negative ionizable groups are attached to silver ions. The reports show that the pH is also a determining factor of the shape, size, production rate, and stability of nanoparticles [14,27].

**Figure 4:**
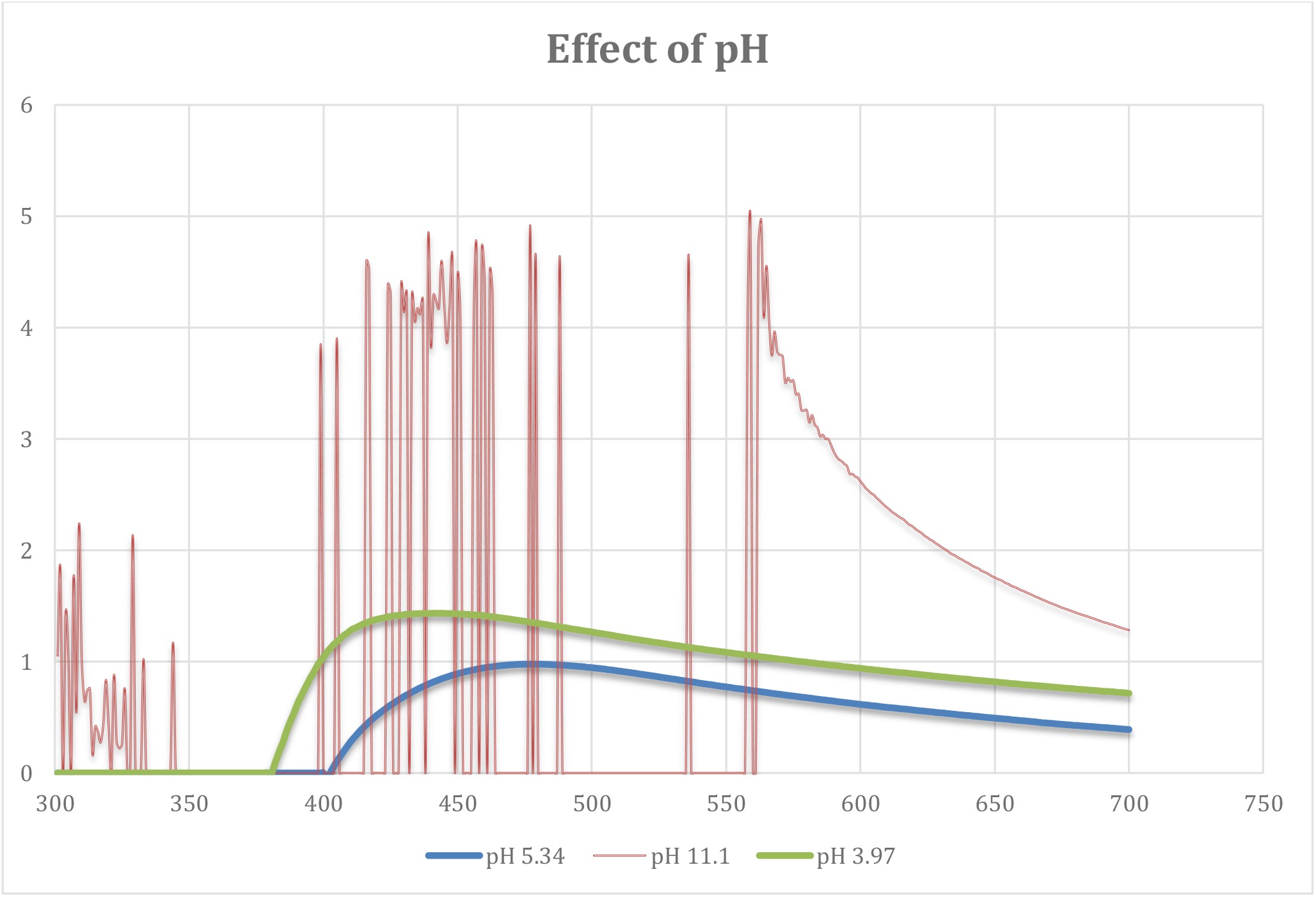
Absorbance spectrum of silver nanoparticles at different pH. For a pH of 3.97, the peak was observed at 435 nm. At a pH of 11.1, a random absorbance pattern is observed.

The concentration of plant extract had a significant impact on the reaction time and the concentration of nanoparticles as we can see from Figure 5. As the amount of plant extract was gradually increased from 8 ml to 12 ml, we could see a significant difference between the reaction medium and the absorbance of the silver nanoparticles. It can be noted that increasing the volume made the reaction rate high with increasing concentration of nanoparticles compared to our standard 8 ml of plant extract. This means that the synthesis of silver nanoparticles is greatly affected by the concentration of reducing and stabilizing precursors which in this case is the plant extract used in the reaction. The increased number of biomolecules results in agglomeration, which reduces the absorption in UV-Vis spectroscopy[14]. In general, the particle size and the reaction rate can be controlled by optimizing the amount of plant extract used in the reduction process.

**Figure 5:**
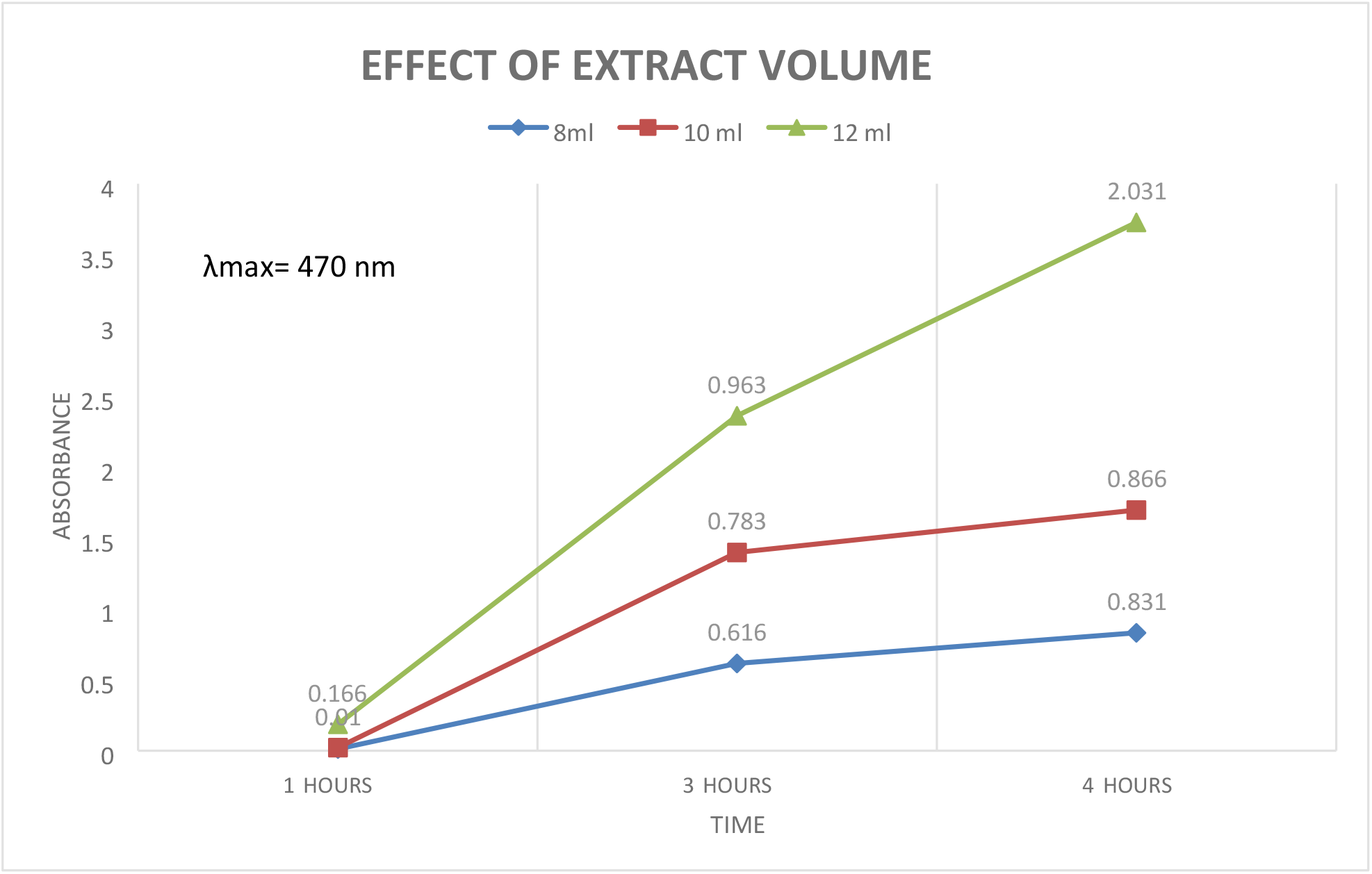
Effect of Plant extracts volume on the reaction time. At 3-hours, the sample containing the highest quantity of plant extract showed the highest absorbance, recorded at 0.963 at λ_max_ calculated to be 470 nm. The lowest absorbance, 0.616, was shown by the lowest volume of plant extract, 8 ml. At the 4-hour mark, the highest was for, once again, the 12 ml of plant extract, showing absorbance of 2.031. The lowest point observed was for the 8 ml plant extract showing absorbance of 0.831.

To determine the particle size in the solutions, we performed several calculations with the computer simulation program Mie Plot v. 4.6.13. It is based on Mie scattering intensity theory. As one can see, in the absorbance spectra, there is only one peak, so we can conclude that our particles are spherical (Figure 6). We can determine the size of particles from the absorbance spectra compared with the calculated one using Mie theory. The peak position directly depends on the size of nanoparticles, as the size of the particle increases, the SPR position shifts to longer wavelengths. When the particle size is considered 47 nm, the spectrum we get from the theoretical model is the closest match to our experimental values. As we can see from the theoretical model that there is a presence of a secondary peak, which means it has dipole Plasmon resonance, and there are two different sizes of nanoparticles present in the medium. The theoretical spectrum has a higher absorbance than our experimental curve. This could be since we have performed the experiment on a smaller scale. There is a limited number of particles present in the medium, hence the lower absorbance value than our reference model. However, these calculations are not exact and cannot confirm that the nanoparticles’ size is the actual size we synthesized through the process. Instead, they give us a rough estimate about the silver nanoparticles’ size and shape[28]. These calculations have another disadvantage, and that is, it can be applied only in spherical particles. The only way to accurately determine the size and shape of the nanoparticles is through FTIR analysis of the silver nanoparticles[5].

**Figure 6:**
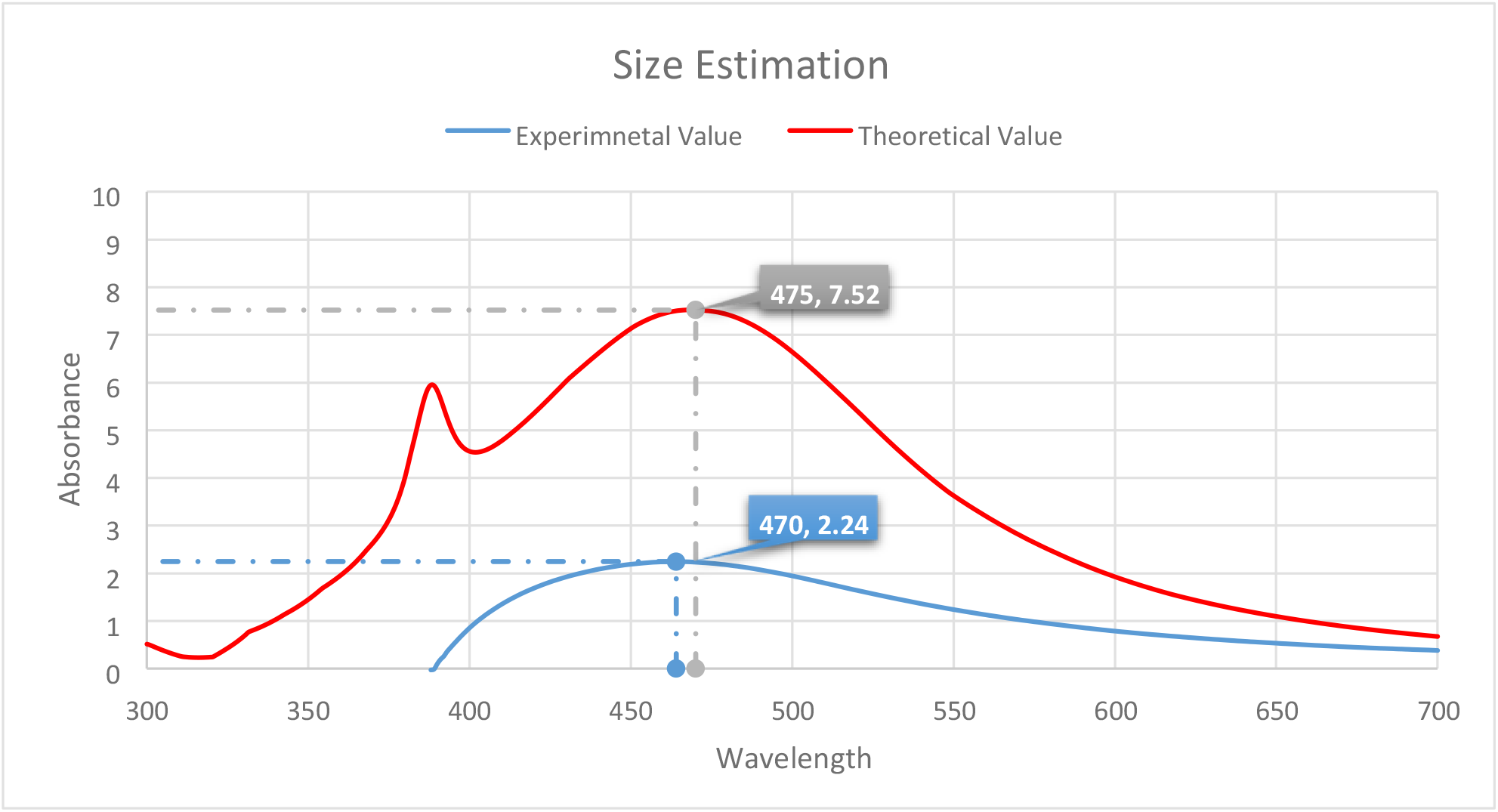
Estimating the size of the Silver Nanoparticles. For the experimental value, the absorbance was revealed to be 2.24 at a wavelength of 470. This is compared with the theoretical value, with an absorbance of 7.52, and a wavelength of 475, to estimate the size of the silver nanoparticles formed.

One of the most crucial aspects of the nanoparticle should be its stability. This means the particles must remain stable either in the suspension or in their solid powder form. If the particles are not stable, it would be impossible to use them for anything as they would be challenging to store under normal conditions. When the nanoparticles are synthesized through physical or chemical means, stabilizers are added to keep them viable for more extended periods. However, plant extract mediated synthesis of nanoparticles does not require stabilizers as the plant phytochemicals keep them stable and maintain the shape and size of the nanoparticles. In our case, the silver nanoparticles were stable for four days while kept at room temperature (Figure 7). Sunlight can significantly influence the size and shape of the nanoparticles. Sunlight can cause auto-oxidation of the nanoparticles and destabilizes them. Usually, these particles are kept away from sunlight and stored in a dark room. In-plant extract mediated synthesis, the sunlight cannot harm the particles as they are surrounded by plant phytochemicals. The curves for each time frame are similar to each other, which means that the shape of the nanoparticles was unchanged through this period (Figure 8). Then again, the SPR values remained constant, meaning that the particles could retain their shape without the stabilizers’ addition.

**Figure 7:**
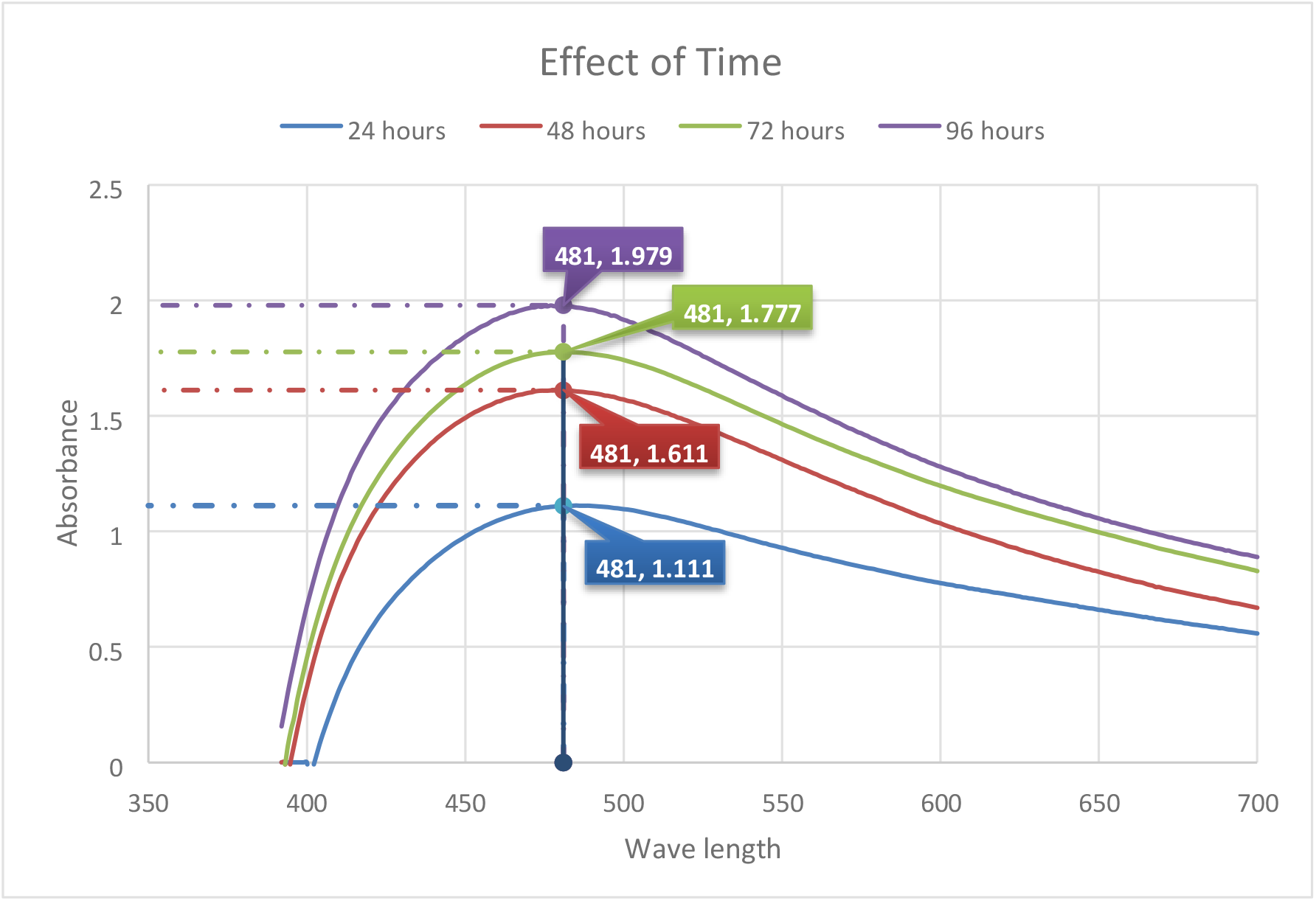
Stability of the Synthesized Silver Nanoparticles. The absorbance peaked at 1.111, with the SPR positioned at 481 nm at 24 hours. The maximum absorbance was observed at the 96-hour mark showing absorbance of 1.979. At the 48 hour mark, the absorbance was 1.611 at an SPR of 481 while at the 72 hour mark, the absorbance was 1.777 at an SPR of 481. Little to no absorbance was observed for all of them below 390 nm.

**Figure 8:**
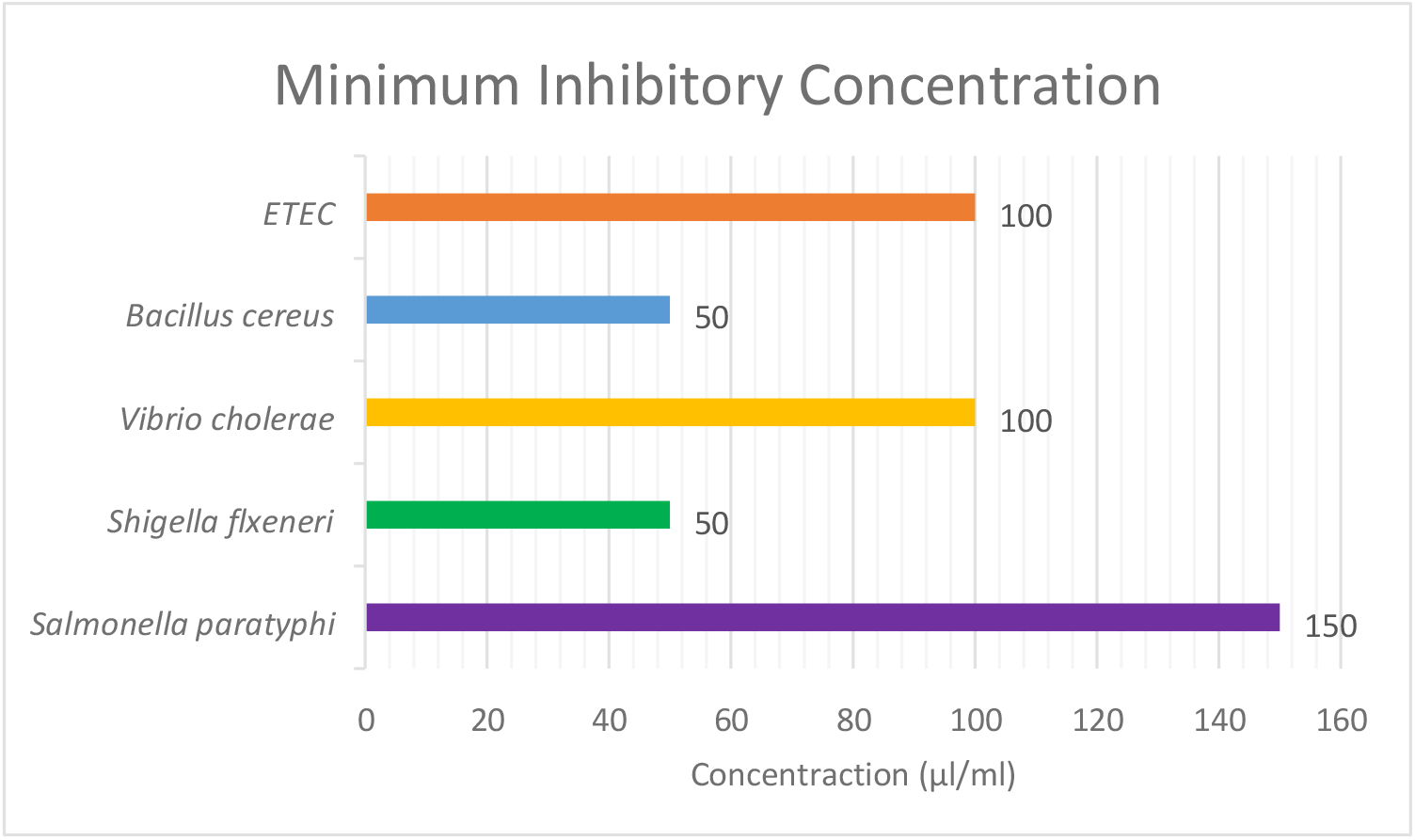
Results of the Minimum inhibitory test. *Salmonella paratyphi* required the highest concentration of silver nanoparticles at 150 µl/ml while the lowest amount, 50 µl/ml, was required by *Bacillus cereus* and *Shigella flexneri*. ETEC as well as *Vibrio Cholerae* required the same concentration of silver nanoparticles at 100 µl/ml.

One of the main objectives of this research project was to check the activity of nanoparticles against different types of pathogenic organisms. The results of the antimicrobial assay have been described in Table 1. The literature review discussed that silver nanoparticles could show enhanced antimicrobial activity when combined with the plant extract. Our results indicate that there is undoubtedly an increase in the combined therapy inhibition zones, but it wasn’t significant enough since the rise is too small. From Table 1, we can see that in the case of *E. coli* (ETEC), the nanoparticles showed an inhibition zone of 11.5 mm, but when combined with plant extract, the inhibition zone slightly increases to 12 mm, which is not much to say anything for sure. This could’ve happened for several different reasons.

**Table 1:** Zone of inhibition of different organisms (all measurements indicated are in mm). The antibacterial assay shows that the largest zone of inhibition was observed for *Vibrio Cholerae* in the case of plant extract and Nanoparticles, while the lowest zone of inhibition was seen in the case of *Bacillus Cereus* incase of Nanoparticles. Results of the Agar Well Diffusion for Nanoparticles showed that the largest zone of inhibition was given by *Vibrio Cholerae* while the lowest was achieved by *Bacillus Cereus*. The zone of inhibition was zero for all pathogens in the Plant extract. The difference between the size of zone of inhibitions for plant extract and nanoparticles alone, compared to a combination of plant extracts and nanoparticles is not significant.

| Organism | AgNO<sub>3</sub> | Plant extract | Nanoparticles | Plant extract and Nanoparticles |
| --- | --- | --- | --- | --- |
| *Salmonella Paratyphi* | 11 | 0 | 11.5 | 12 |
| *Shigella Flexneri* | 11 | 0 | 12 | 13 |
| *Vibrio Cholerae* | 13 | 0 | 13 | 14 |
| *Bacillus cereus* | 11 | 0 | 10 | 11 |
| *E.coli (ETEC)* | 11.5 | 0 | 11.5 | 12 |

First, the purification process of the nanoparticles wasn’t performed according to the already established protocol. This was due to the lack of proper equipment while experimenting. As a result, repeated centrifugation was needed, which can sometimes destabilize the particles and significantly hamper their antimicrobial properties. Another important reason could be that the plant extract concentration used in this process was deficient compared to our standard protocol. This was done because the plant extract concentration can affect the shape and optical properties of the silver nanoparticles. Optimization is needed in the extract concentration to have a perfect balance between the nanoparticle synthesis and its ability to show antimicrobial activity. Increasing the plant extract concentration might yield better results and thus increase the inhibition zones.

## 4.2 Conclusion

To conclude, we can say that the objectives set forth before starting this research project have been successful. The primary object was to determine whether the aqueous extract of *Cymbopogon citratus* can synthesize silver nanoparticles when AgNO_3_ is used as a precursor molecule. The results of the UV spectral analysis clearly show the formation of nanoparticles in the solution. This research was not aimed towards using the silver nanoparticles as a substitute for our conventional antibiotics. Rather, it was a steppingstone to show that we can synthesize nanoparticles without using toxic components. Cytotoxic analysis can reveal the adverse effect the nanoparticles have upon consumption by the test, giving us an idea about the dosage requirements without harming the host cell components. Nanoparticles synthesized through chemical means have tumor-suppressing capabilities. Since, our nanoparticles show almost similar optical properties, it can also be tested against cancer cells to see if they can inhibit their growth and potentially kill them. Moreover, green synthesis of nanoparticles can clean the environment and be used as biosensors to detect contamination. Most importantly, the process used to create the particles doesn’t require any toxic compounds, which is one of the main drawbacks of chemical methods. The field of Nanotechnology is growing and still developing.

The findings of this research project can influence further studies in this field to understand the technology better and gain more knowledge about the properties and applications of nanoparticles.

